# Replication of 10 novel loci involved in human plasma protein N-glycosylation using MALDI-MS and UHPLC-FD data

**DOI:** 10.1101/2025.08.01.668131

**Authors:** Anna Timoshchuk, Annemieke Naber, Roderick Slieker, Anna Soplenkova, Denis Maslov, Nadezhda A. Potapova, Simone Nicolardi, P.J.M. Elders, Eric J.G. Sijbrands, Sodbo Sharapov, Leen M. ‘t Hart, Mandy van Hoek, Manfred Wuhrer, Yurii S. Aulchenko

**Author notes:** Correspondence to: Yurii S. Aulchenko. These authors contributed equally (Annemieke Naber, Roderick Slieker, Anna Soplenkova, Denis Maslov, Nadezhda A. Potapova). Current address: GSK Medicines Research Centre; Gunnels Wood Road, Stevenage, SG1 2NY, UK.

## Abstract

N-glycans are essential components of glycoproteins, influencing their properties and functions. While biochemical pathways of glycosylation are well-characterized, their genetic regulation remains poorly understood. This study utilizes matrix-assisted laser desorption/ionization-mass spectrometry (MALDI-MS) and ultra-high performance liquid chromatography-fluorescence detection (UHPLC-FD) to strengthen replication and further characterize previously identified genome-wide association signals for the total human plasma N-glycome (TPNG). Univariate and multivariate genetic association meta-analyses involved 3,385 samples across 143 N-glycome traits from the Hoorn Diabetes Care System and DiaGene cohorts as well as 3,224 samples across 117 N-glycome traits from TwinsUK, CEDAR, QMDiab and SABRE cohorts. We successfully replicated ten previously identified but not replicated glycosylation quantitative trait loci (glyQTLs) and prioritized five high-confidence putative causal genes, including the glycosyltransferase *MGAT4B* and inflammation-related genes – *C3* and *FCGR2B.* The linkage-specific sialic acid derivatization in MALDI-MS enabled delineation of genetic effects on α2,3- and α2,6-sialylation. Mass spectrometry analysis also provided evidence for glucuronic acid-containing glycans in human blood plasma. These findings advance our understanding of the genetic regulation of protein N-glycosylation and highlight the complementarity of different analytical approaches in glycomics research.

## Introduction

N-glycans are irregular polymers attached to polypeptide structures, forming glycoproteins with a glycosidic bond to the asparagine (Asn) side chain (1). Glycosylation impacts the physical and chemical characteristics of proteins, along with their biological roles. Changes in protein glycosylation are associated with a number of human diseases, and glycans are seen as biomarkers and essential components of treatments (2,3). Although the biochemical pathways of glycosylation are well understood (4), genetic and regulatory networks controlling glycan diversity, tissue-specific expression, and disease-related variation remain poorly understood. The lack of understanding of mechanisms governing the variation in the glycome and connecting it to human health and disease delays the progress in the clinical application of human glycobiology (2,3).

In the most recent genome-wide association study (GWAS) of total plasma N-glycome (TPNG), we used an ultra-high performance liquid chromatography-fluorescence detection (UHPLC-FD) to measure N-glycans in about 10,000 analyzed samples (5). This study discovered 59 TPNG glycan quantitative trait loci (glyQTLs), 40 of which were replicated in 3224 independent samples.

Understanding the spectrum of glycans associated with a specific locus is crucial for interpreting the association of the locus, including prioritization of candidate genes and formulating hypotheses about mechanisms through which genetic variation in the locus affects the phenotype. For instance, when a locus contains a glycosyltransferase gene, and the variation associated with this locus concerns N-glycans that are expected to be affected by the glycosyltransferase, this strongly supports the glycosyltransferase as the primary candidate gene in that region. Examples include association of the *FUT8*-containing region with core-fucosylated N-glycans and their substrates, and the *MGAT3* region with bisecting GlcNAc (6). Phenotypic variation can indicate the tissue of action of the gene, e.g. a locus affecting tri- and tetra-antennary structures of TPNG is probably acting in the liver, as these structures are predominantly expressed on liver-secreted glycoproteins (6).

Conversely, a mismatch between expected action of a candidate gene and a strong biological prior (such as a glycosyltransferase) and the phenotypic effects of the locus may suggest an alternative candidate gene, a complex, e.g., indirect, mechanism of action, or a methodological artifact. For instance, the *B3GAT1* gene, which encodes a key enzyme involved in the glucuronyl transfer reaction, is a candidate for being a causal gene in one of the first loci linked to TPNG (7). However, glucuronic acid monomers are not typically found in the N-glycome of human blood plasma. These associations triggered mass spectrometric analysis of the associated TPNG peaks which confirmed the presence of glucuronic acid-containing N-glycans (7). Another example concerns, the ABO locus that encodes glycosyltransferase enzymes known to convert H antigen substrates into blood group antigens. However, plasma glycoproteins lack the H antigen substrate, making it unlikely that the genetic association between the *ABO* locus and TPNG is driven directly by *ABO* glycosyltransferase activity, suggesting instead an indirect or alternative mechanistic explanation.

Utilizing different analytical techniques allows for a more detailed understanding of the glycan spectrum associated with specific loci, supporting the goal of deciphering the biology of associated loci. Most previous studies of quantitative genetics of glycans employed HPLC (6), while other high-throughput technologies for TPNG exist (8). These include multiplexed capillary gel electrophoresis with laser-induced fluorescence detection (xCGE-LIF) (9), and matrix-assisted laser desorption ionization mass spectrometry (MALDI-MS). UHPLC-FD and xCGE-LIF technologies allow separating structural isomers of N-glycans, providing branch-specific information, that is, a separation between the 3-arm and 6-arm isomers of glycan species (for example, FA2[3]G1 and FA2[6]G1). MALDI-MS is unable to differentiate the structural isomers.

In the current study, we took advantage of analyzing data generated by UHPLC-FD and MALDI-MS. One of the unique features of the latter method is linkage-specific separation of α2,3- and α2,6-sialylation due to specific sample preprocessing. This distinction is particularly valuable since previous studies demonstrated that, these two sialylation types may have different, and even opposite associations with diseases. For example, diabetes and liver fibrosis in metabolic dysfunction-associated steatotic liver disease (MASLD) are associated with higher α2,6-sialylation and lower α2,3-sialylation (10,11).

In the present study, our objectives were to replicate the association of the 19 loci, previously reported to be associated with TPNG at GWAS significance level, but not yet replicated (5,12), and to prioritize genes, likely playing a role in regulation of N-glycosylation of plasma proteins, in newly replicated loci. Further, we aimed to utilize UHPLC-FD and MALDI-MS N-glycan spectra to provide extended annotation of the phenotypic effects, i.e., associated N-glycans, of the loci with confirmed association with TPNG (i.e. replicated here and elsewhere) (5,12).

## Results

### Replication

Two groups of cohorts were used for replication. The first one involved 3,224 samples from several cohorts phenotyped with UHPLC-FD across 117 N-glycome traits, that were used for replication in Sharapov et al. (5). The second group is unique to this study and includes 3,385 samples from the Hoorn Diabetes Care System and DiaGene cohorts, phenotyped for 143 N-glycome traits with MALDI-MS. A per-trait, per-locus meta-analysis was applied separately to cohorts within the two groups. As we cannot fully match the N-glycan traits measured by UHPLC-FD and MALDI-MS, for each of the 19 glyQTLs not replicated in Sharapov et al., we applied the Cauchy aggregation test (13) to combine P-values across all glycan traits, resulting in a single p-value per locus per group. We then conducted a Fisher product meta-analysis (14) for Cauchy aggregated p-values from UHPLC-FD and MALDI-MS groups, resulting in 19 replication p-values (Supplementary Table 3a). A locus was considered replicated if the Fisher product meta-analysis p-value exceeded the threshold of 0.05/19=2.63E-03, where 19 is the number of glyQTLs analyzed here.

We replicated 10 loci (containing *FCGR2B, NDUFB4P4, PCCB, RNF168, MGAT4B, GRID1P, ODF1, OVOL1, RPLP2P4, C3* genes) out of 19 tested for replication (Table 1). The details of the replication procedure are presented in the Supplementary Table 3b.

**Table 1.**
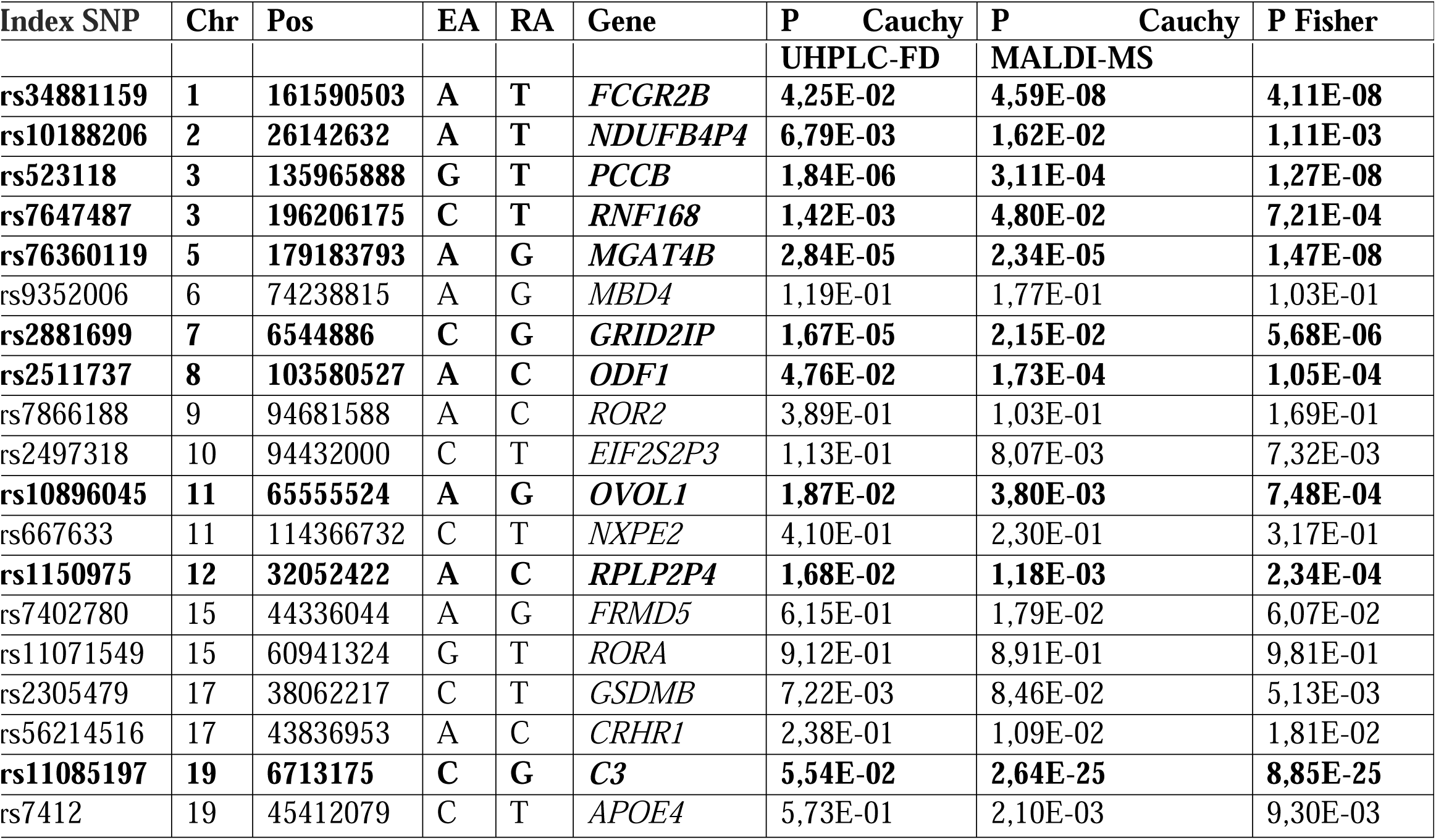
Results of the replication of 19 loci not replicated in the study by Sharapov, et al. (5). Ten loci (in bold) are replicated in the current study with a threshold of the Fisher product meta-analysis p-value of 2.63E-03. Full results are presented in Supplementary Table 3a. Chr— chromosome; Pos—position on chromosome according to GRCh37.p13 assembly; EA—Effect allele; RA—Reference allele; Gene—Nearest gene or gene prioritized in this work or in previous GWAS for human plasma protein N-glycome; P Cauchy UHPLC-FD—Cauchy aggregated P-value across all UHPLC-FD-measured glycan traits; P Cauchy MALDI-MS—Cauchy aggregated P-value across all MALDI-MS-measured glycan traits; P Fisher—Fisher product meta-analysis P-value.

### Prioritization of causal genes for protein N-glycosylation

We used several approaches to prioritize the most likely effector genes for the ten loci replicated in this work: prioritization of genes encoding glycosyltransferases; genes causing congenital disorders of glycosylation (CDG); colocalization of glyQTLs with gene expression QTLs (eQTLs) in liver and whole blood, and loci associated with blood plasma protein levels (pQTLs); annotation of putative causal variants affecting protein structure; enrichment of gene sets and tissue-specific expression; and prioritization of the nearest gene (see Methods).

In total, 22 candidate genes showed at least one indication for prioritization in 10 replicated glyQTLs (Figure 1a, Supplementary Table 4a). Among these, we identified two genes in the same locus encoding glycosyltransferases (*MGAT5B, MGAT4B*). The Summary data-based Mendelian Randomization analysis followed by the Heterogeneity in Dependent Instruments (SMR/HEIDI) approach (15) indicated that total plasma N-glycosylation–associated variants in two loci possibly had pleiotropic effects on transcription of three genes (*FCGR2B*, *FCGRLB, NCK1*) in relevant tissues (Supplementary Table 4f). In 3 genes (*RNF168*, *MGAT4B*, *C3*), associated variants were either coding or were in strong LD with the variants coding for potentially deleterious amino acid changes (annotated by Variant Effect Predictor, VEP) (16). The DEPICT gene prioritization tool (17) provided evidence of prioritization for 12 genes in 6 loci at FDR < 0.2 (Supplementary Table 6h). We didn’t prioritize any genes based on literature annotations related to CDG genes, and none of genes showed colocalization with whole blood pQTLs.

**Figure 1.**
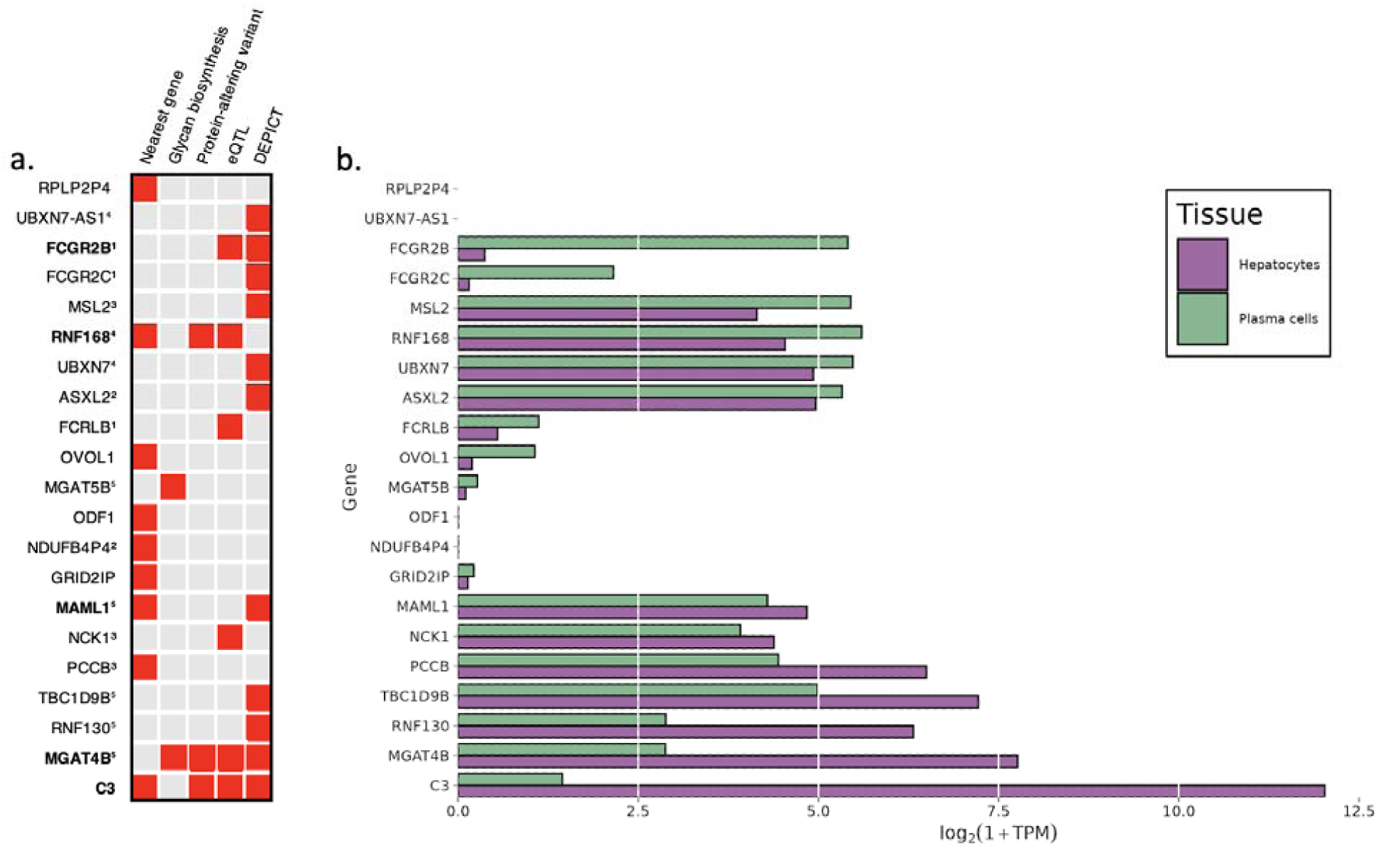
Candidate genes. (a) Predictors indicating 22 candidate genes. The identical superscripts denote candidate genes inside one locus. Five high-confidence putative causal gene are highlighted in bold. Full details of the gene prioritization are presented in Supplementary Table 4a. (b) Gene expression of the candidate genes in two relevant cell types: hepatocytes and plasma cells. Expression levels are represented as the median logarithm of transcripts per million. The data for hepatocytes (N = 513) and plasma cells (N = 53) samples were obtained from the ARCHS4 portal (19).

We required a candidate gene to be expressed in liver or plasma cells, because these two tissues are the main producents of N-glycosylation (18) (Figure 1b). We prioritized it as potentially causal if at least two pieces of evidence supported it. This resulted in prioritizing five high-confidence putative causal genes: *FCGR2B, RNF168, MAML1, MGAT4B,* and *C3*), highlighted in bold in Figure 1a, which will be the focus of our subsequent discussion.

### Characterization of glyQTLs by spectrum of associated N-glycan traits

We compared annotations, specifically R^2^ of association between a locus and a top trait, for all glycosyltransferase loci (Supplementary Tables 6a, 6b). We observed a consistency between associated spectra of glycans measured with UHPLC-FD and MALDI-MS. Specifically, the *FUT8* locus (fucosyltransferase 8, an enzyme responsible for the addition of core fucose to glycans) was associated mostly with core-fucosylated di-antennary glycans and tri- and tetra-antennary fucosylated glycans. The *FUT6* locus showed associations with a spectrum of tri- and tetra-antennary fucosylated sialylated glycans. The *MGAT3* (beta-1,4-mannosyl-glycoprotein 4-beta-N-acetylglucosaminyltransferase) locus was associated with bisected glycans, reflecting its enzymatic activity. The locus containing *MGAT5* gene that catalyzes the addition of beta-1,6-N-acetylglucosamine to the alpha6-linked mannose of di-antennary glycans showed associations with a variety of tri- and tetra-antennary sialylated structures. The *B4GALT1* gene encodes galactosyltransferase, which adds galactose during the biosynthesis of different glycoconjugates. This locus was associated with galactosylation of di-antennary glycans. Annotation of other loci can be found in Supplementary Tables 6a, 6b.

After that, we annotated the loci that had the potential to be uniquely characterized by MALDI-MS: *ST6GAL1*, *ST3GAL6*, *ST3GAL4* and *B3GAT1* (Supplementary Table 6b).

By derivatizing sialic acid with linkage-specific ethylation and lactonization, we could distinguish between α2,6-linked and α2,3-linked sialic acid residues based on their mass. In order to identify the benefits of sialic acid derivatizing provided by MALDI-MS compared to UHPLC-FD we used the following three benchmarking loci: *ST6GAL1*, *ST3GAL6*, *ST3GAL4* (Table 2). The locus containing the *ST6GAL1* gene (beta-galactoside alpha-2,6-sialyltransferase 1) was linked to a proportion of α2,6-sialylated galactosylated di-antennary glycans and several derived traits that predominantly combine sialylated and non-sialylated galactosylated di-antennary glycans (top R^2^=0,019 with directly measured trait H4N4F1E1 which corresponds to a di-antennary core-fucosylated α2,6-sialylated glycan). The locus for the *ST3GAL6* gene (beta-galactoside alpha-2,3-sialyltransferase 6) demonstrated an association with a proportion of α2,3-sialylated galactosylated tetra-antennary glycans (top R^2^= 0,009 with derived trait A4L representing α2,3-sialylation per antenna within tetra-antennary glycans). Additionally, the *ST3GAL4* locus (beta-galactoside alpha-2,3-sialyltransferase 4) was associated with a wide range of sialylated glycans, primarily α2,3-sialylated, including di-, tri-, and tetra-antennary structures (top R^2^= 0,140 with derived trait A3L which corresponds α2,3-sialylation per antenna within tri-antennary glycans).

**Table 2.** Annotation of the sialyltransferase loci. Full annotation results are presented in Supplementary Tables 6a, 6b. For R^2^ calculation, we used SNPs that showed the most significant association with N-glycome trait from (5).

| Glycan Trait (MALDI-MS) | Corresponding glycan trait (UHPLC-FD) | Measurement technology | $R^2 \times 100$ of association with *ST3GAL6* | $R^2 \times 100$ of association with *ST3GAL4* | $R^2 \times 100$ of association with *ST6GAL1* |
| --- | --- | --- | --- | --- | --- |
| sialylation per antenna within all complex glycans | the percentage of sialylated structures in total plasma glycans | UHPLC-FD | 0,00 | 0,55 | 0,02 |
|  |  | MALDI-MS | 0,00 | 0,11 | 0,03 |
| α2,6-sialylation per antenna within diantennary glycans | the percentage of disialylated structures in total plasma glycans | UHPLC-FD | 0,00 | 0,49 | 0,01 |
|  |  | MALDI-MS | 0,05 | 0,18 | 0,00 |
| α2,6-sialylation per antenna within triantennary glycans | the percentage of trisialylated structures in total plasma glycans | UHPLC-FD | 0,01 | 0,04 | 0,16 |
|  |  | MALDI-MS | 0,45 | 7,82 | 0,15 |
| α2,6-sialylation per antenna within tetraantennary glycans | the percentage of tetrasialylated structures in total plasma glycans | UHPLC-FD | 0,10 | 1,05 | 0,02 |
|  |  | MALDI-MS | 0,61 | 8,76 | 0,18 |
| α2,3-sialylation per antenna within diantennary glycans | the percentage of disialylated structures in total plasma glycans | UHPLC-FD | 0,00 | 0,49 | 0,01 |
|  |  | MALDI-MS | 0,23 | 6,80 | 0,01 |
| α2,3-sialylation per antenna within triantennary glycans | the percentage of trisialylated structures in total plasma glycans | UHPLC-FD | 0,01 | 0,04 | 0,16 |
|  |  | MALDI-MS | 0,64 | 13,99 | 0,07 |
| α2,3-sialylation per antenna within tetraantennary glycans | the percentage of tetrasialylated structures in total plasma glycans | UHPLC-FD | 0,10 | 1,05 | 0,02 |
|  |  | MALDI-MS | 0,92 | 12,17 | 0,11 |

Interestingly, on average the strength of genetic association between loci containing sialyltransferases and MALDI-MS-measured N-glycome traits was 5.4 times higher, when compared with UHPLC-FD-measured N-glycome traits. Multiple factors or their combination can explain this difference. Firstly, the noise-attributed variance component of MALDI-MS measured traits can be lower than that in UHPLC-FD. Second, the current set of derived traits differs between MALDI-MS and UHPLC-FD, and MALDI-MS-derived traits may better reflect the enzymatic activity of sialyltransferases.

The locus containing the *B3GAT1* gene encoding beta-1,3-glucuronyltransferase showed the most significant association with H5N5E1 glycan (R^2^=0.043). In previous studies, glucuronic acid was not detected in the TPNG profile, but only in glycan profiles obtained after TPNG desialylation (7). Here, we analyzed the H5N5E1 (Figure 3A) glycan peak and identified an overlapping, glucuronic acid-containing species in the blood plasma glycome. Ultrahigh resolution MALDI-MS measurements of TPNG profiles detected the bisected, 2,6-sialylated H5N5E1 glycan (Figure 3A) at *m/z* 2185.7882 with high accuracy. The partially overlapping species detected at *m/z* 2186.7713 matched with a composition of H5N4E1G1, where G1 stands for a hexuronic acid moiety that underwent ethyl esterification during the sialic acid derivatization step (Figure 3B). This independent confirmation of glucuronic acid-containing species in native TPNG profiles strengthens the notion of their presence in human blood plasma, warranting further investigation.

**Figure 3.**
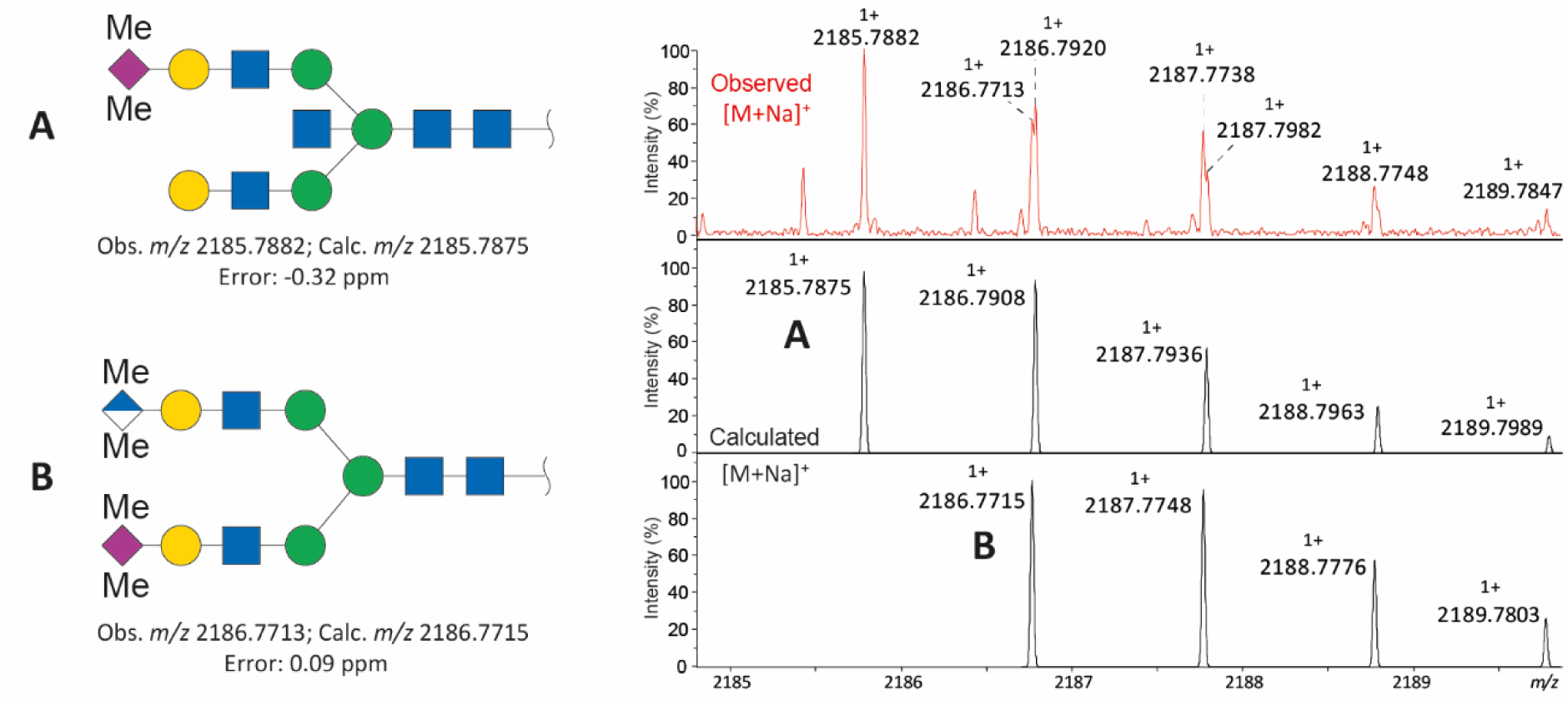
Ultrahigh resolution MALDI-FT-ICR-MS supports the coexistence and partial overlap of the isotopic peaks of the glucuronic acid-containing species and a bisected, sialylated N-glycan. **(A).** The top panel shows a representative measured spectrum. The lower two panels show the calculated spectra of species A and B. Glycan color code: blue square, N-acetylglucosamine; green circle, mannose; yellow circle, galactose; purple diamond, sialic acid; blue/white diamond, glucuronic acid; Me, methylation.

## Discussion

In this study, we replicated, for the first time, ten previously reported TPNG GWAS loci - containing *FCGR2B, NDUFB4P4, PCCB, RNF168, MGAT4B, GRID1P, ODF1, OVOL1, RPLP2P4, C3*. We prioritized five high-confidence putative causal genes for these loci (*FCGR2B, RNF168, MAML1, MGAT4B, C3*) (Table 3).

**Table 3.**
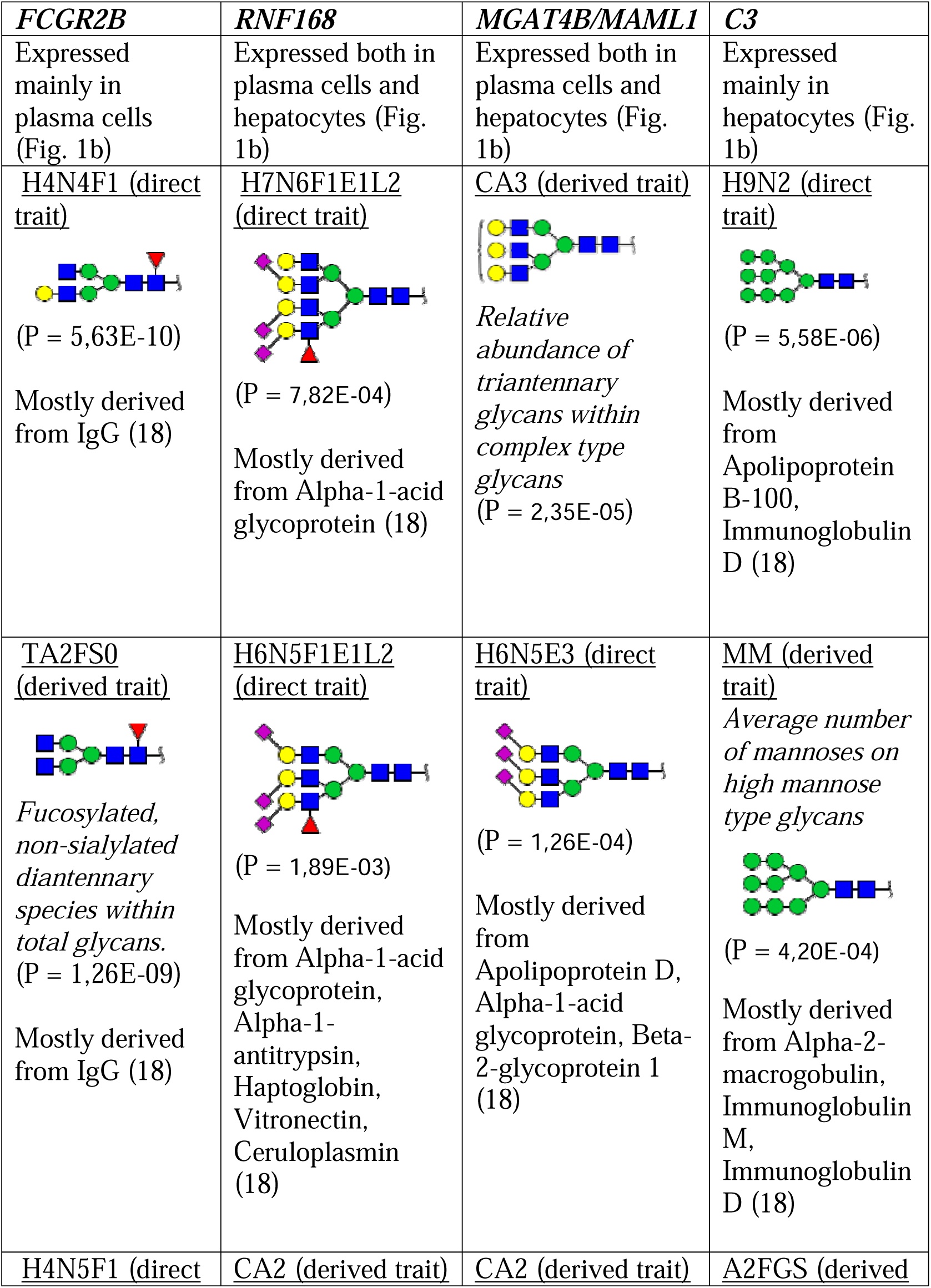
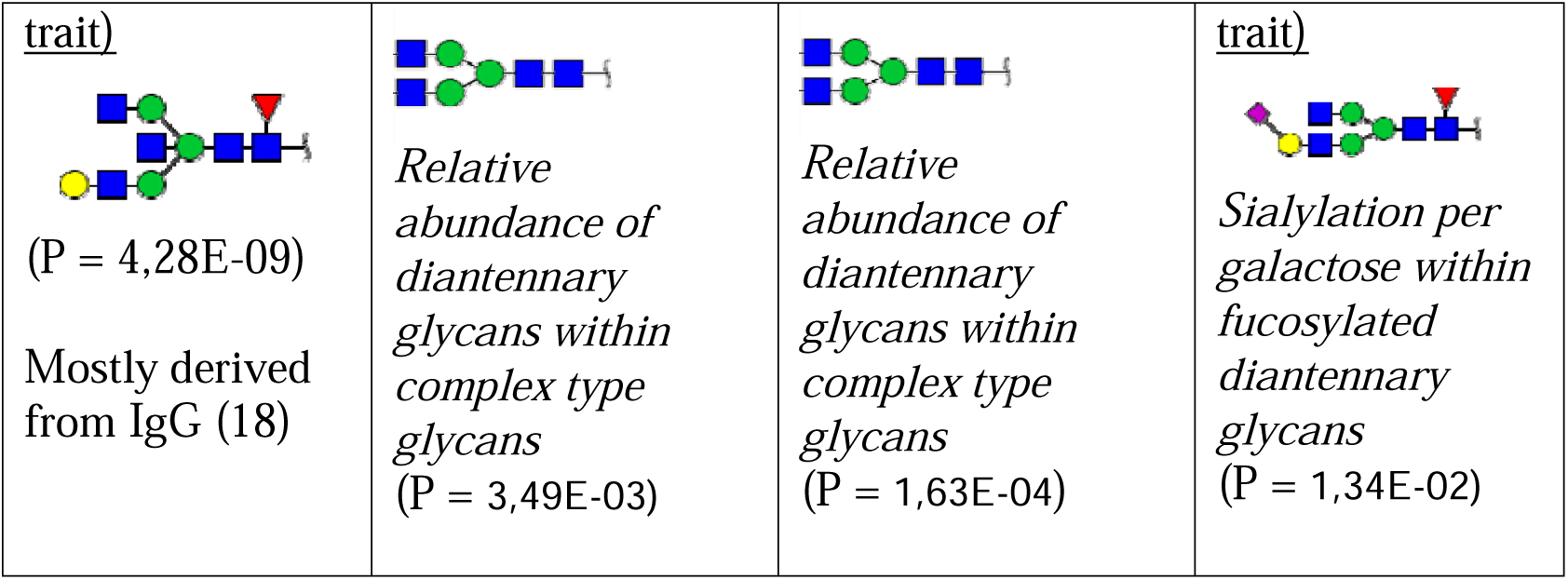
Annotation of four loci containing high-confidence putative causal genes. For each locus, the top three associated N-glycome traits are represented. Full annotation results are presented in Supplementary Table 6b.

The *FCGR2B* gene encodes the Fc gamma receptor IIB (CD32B), which plays a critical role in the immune response by binding to the Fc region of IgG. CD32B is primarily an inhibitory receptor expressed by B cells, basophils, and monocytes. CD32B inhibits phagocytosis and pro-inflammatory cytokine release, primarily by competing with different activating Fc gamma receptors for available immune complexes (20). Additionally, it has been shown that crosslinking of CD32B to the B cell receptor increases the B cell activation threshold (20), thus decreasing antibody production and influencing the N-glycome. Dysregulation in its function could caus increased immune responsiveness and predisposition to autoimmunity. Consistent with this, downregulation of *FCGR2B* expression has been reported in systemic lupus erythematosus (21).

In this locus the *FCRLB* and *FCGR2C* genes are located in close proximity and may also contribute to alterations in glycosylation pattern. *FCGR2C* encodes an activating Fc gamma receptor that is expressed on NK cells and, unlike *FCGR2B*, it promotes immune activation and mediates antibody-dependent cellular cytotoxicity. Interestingly, the gene harbors a well-characterized protein altering polymorphism in the third exon - rs759550223 (*57X/Q*) – which is in linkage disequilibrium with the lead polymorphism for this locus (rs34881159, D’=0.7686, p=0.026). The reference allele (T) introduces a premature stop-codon rendering the receptor non-functional leading to variability in immune responses. This functional variation can influence the plasma N-glycome by modulating immune complex handling and antibody effector functions (22,23).

The *MGAT4B* gene encodes the enzyme N-acetylglucosaminyltransferase IVB (GnT-IVb), which catalyzes the transfer of GlcNAc from UDP-GlcNAc to the GlcNAc-beta1,2-Man-alpha-1,3 arm of the core structure of N-linked glycans through a beta-1,4 linkage. This enzymatic activity is essential for generating of complex-type N-glycans, characterized by their branched structures, which are crucial for various cellular functions, including cell signaling, adhesion, and the modulation of immune responses (24).

The *MAML1* gene, also known as Mastermind-like 1, is a transcriptional co-activator that plays a crucial role in the Notch signaling pathway. This pathway is involved in various cellular processes, including cell fate determination, proliferation, and differentiation. MAML1 interacts with the intracellular domain of Notch receptors and other transcription factors to regulate gene expression (25). In the current work, we have firstly prioritized *MAML1* as a potential regulator of N-glycosylation, proposing that it may regulate the transcription of specific glycosyltransferases through its interaction with Notch. Further research is warranted to figure this out.

The *C3* gene encodes complement protein C3 which plays a central role in the classical, alternative and lectin activation pathways of the complement system. Earlier, we had highlighted the *CFH* (complement factor H) gene as a potential regulator of N-glycosylation (5). It was speculated that genetic variation in *CFH* affects the spectrum of N-glycans attached to immunoglobulins through the regulation of inflammation. CFH is the major inhibitor of the alternate complement pathway, accelerating the decay of alternative C3 convertase, thus inhibiting C3 cleavage, essential for downstream pathway action (26). Considering the compensating effect of these two proteins on the complement factor activity, it is tempting to assume a common mechanism for their effect on the composition of total plasma N-glycome.

The gene *RNF168* (ring finger protein 168), replicated previously in multivariate IgG GWAS (27), encodes an E3 ubiquitin ligase protein involved in DNA double-strand break repair and immunoglobulin class switch recombination (28). Mutations in this gene lead to the very rare RIDDLE syndrome (an immunodeficiency and radiosensitivity disorder), with an estimated prevalence of fewer than 1 in 1,000,000 individuals. The features of this syndrome include decreased immunoglobulin production (29). An association between SNPs at/near *RNF168* and IgM levels was previously reported (30). As far as we are aware, *RNF168* involvement in glycosylation processes has not been reported, but if so, a pleiotropic action on immune-related and glycosylation traits may be assumed.

Here, we first applied a Cauchy aggregation test to combine the results of UHPLC-FD and MALDI-MS GWASs. This approach allows for a meta-analysis of all traits, in contrast to a meta-analysis that only examines fully matched traits.

Analysis of plasma N-glycome using MALDI-MS technology with linkage-specific sialic acid derivatization provided the first genetic evidence for the tissue- and linkage-specific activity of sialyltransferases *in vivo*, with *ST3GAL4* and *ST3GAL6* associating with α2,3-sialylated glycans and *ST6GAL1* with α2,6-sialylated structures. The spectrum of glycans associated with glycosyltransferase loci, including *FUT8*, *FUT6*, *MGAT3*, *MGAT5*, and *B4GALT1*, aligned with their known enzymatic activities. Mass-spectra associated with variation in the *B3GAT1* locus pointing at glucuronic acid-containing glycans in the blood plasma proteome.

To conclude, in the present study, we demonstrate that analysis of plasma N-glycome using MALDI-MS with linkage-specific sialic acid derivatization allows higher resolution and power in analysis of genetic associations with α2,3- and α2,6-sialylation (Table 2). This is especially important given the linkage-specific associations to metabolic and liver disease (10,11). Further, we extended the list of loci significantly and replicably associated with total plasma N-glycosylation by 10 (a more than 20% increase).

The genes prioritized in these loci with high confidence add to our understanding of the major pathways involved in the regulation of plasma protein N-glycosylation, i.e., glycan biochemistry (*MGAT4B*), B cell (*FCGR2B*), and liver (*C3*) biology. The top three associations for *MGAT4B* are complex high-branched N-glycans (CA3, H6N5E3, CA2) that corresponds to the enzymatic activity of this glycosyltransferase (24) (Table 3). It should be noted that the locus containing the *FCGR2B* gene, which is expressed in plasma cells, is associated with N-glycome traits predominantly derived from IgG (e.g., H4N4F1) (Table 3) (18).

Prioritization of *C3* is of particular interest as this continues highlighting the role of the complement system, which has been implicated by the effects of *CFH* on the plasma N-glycome in a recent work of Sharapov et al. (5). *C3* is primarily expressed in hepatocytes, and the locus containing this gene is predominantly associated with high-mannose traits found in liver-derived proteins (Table 3) (6). This association supports the regulation of N-glycosylation through liver biology. Deeper understanding of biological pathways involved in regulation of N-glycosylation through high-confidence genetic discoveries will facilitate the use of glycans and glycosylation pathways as therapeutic targets and biomarkers of human diseases.

## Materials and methods

### Study cohort description

This work is based on analyzing of the glycomic and genetic data from two cohorts — the DiaGene study (31) and the Hoorn Diabetes Care System (32).

### DiaGene study

The DiaGene study is a multi-center case-control study of type 2 diabetes in the first and second line of care in the region of Eindhoven, the Netherlands. The aim of the DiaGene study was to unravel the etiology of type 2 diabetes and its complications. The DiaGene study includes 1,886 patients with type 2 diabetes and 854 controls without diabetes, of these, in 1,490 cases and 544 controls, the total plasma N-glycome and genotypes were available for analysis. Virtually all patients with type 2 diabetes living in Eindhoven were approached for inclusion, between 2006 and 2011. The control group consisted of people without any kind of diabetes, Cushing’s disease or metformin use; aged 55 years or older (31).

### Hoorn Diabetes Care System

The Hoorn Diabetes Care System (Hoorn DCS) cohort consists of persons with T2D in primary care from the West-Friesland region of the Netherlands. Enrolment in the cohort started in 1998 and this prospective dynamic cohort currently holds 12,673 persons with T2D, of these, in 1,351 cases, the total plasma N-glycome and genotypes were available for analysis (32).

### Total plasma N-glycome profiling

In total, 2,651 DiaGene study samples (1,490 T2D cases and 544 controls) and 1,351 samples (T2D cases) from the Hoorn Diabetes Care System cohort passed QC. Total plasma N-glycome analysis and mass spectrometry data processing for the DiaGene study was based on the workflow from Reiding et al. (33,34) and is described by Dotz et al. (35). In short, total plasma N-glycome was measured by MALDI-time-of-flight (TOF)-MS on a Bruker ultrafleXtreme instrument after enzymatic glycan release from plasma glycoproteins and linkage-specific sialic acid derivatization (36). Quality control of the mass-spectra and glycan analytes was performed, and samples were excluded in case of low intensity or interferences. In total, 73 direct traits passed quality control, and 91 derived traits were calculated by normalization to their sum for the DiaGene cohort; 68 direct and 86 derived traits were calculated for the Hoorn Diabetes Care System cohort. In the DiaGene cohort, batch correction was performed, based on preparation day, MALDI plate, row and column (5). The traits were harmonized between two datasets (see Supplementary Table 1a). MALDI-FT-ICR-MS was performed on a representative sample as described previously (36).

### Genotyping

#### DiaGene study

The DiaGene study participants were genotyped using the Illumina Global Screening array (GSA v1) and genotypes were imputed to the Haplotype Reference Consortium (HRC) reference panel (r1.1) (37) using the Michigan Imputation Server (38).

### Hoorn Diabetes Care System

In total 1,800 Hoorn DCS samples are genotyped and have TPNG measured. All 1,800 samples are of western European descent. 1,600 T2D patients were genotyped using Illumina Core Exome SNP-array. 200 T2D-controls were genotyped using Affymetrix Axiom SNP-array. All genotypes were HRC imputed.

### Genetic association analysis

#### UHPLC-FD GWAS

We reused summary statistics from the replication cohort (TwinsUK, CEDAR, QMDiab and SABRE) (N=3,224) of univariate and multivariate genetic association analysis and meta-analysis of the association between 117 univariate and 21 multivariate total plasma N-glycome traits and the 19 sentinel variants tagging loci that have been reported at genome-wide significance, but did not replicate in the study of Sharapov et al. (5).

### MALDI-MS GWAS

The genetic association analysis in each cohort was conducted using the same protocol as in Sharapov et al. (5). Briefly, prior to GWAS, the total plasma N-glycome traits were adjusted for sex and age, and the residuals were quantile transformed to normal distribution. We assumed an additive model of genetic effects. GWAS were based on the genotypes imputed from Haplotype Reference Consortium Results (39) or 1000 Genomes project (40). Results of univariate GWAS in DiaGene study and Hoorn DCS cohorts passed a strict quality control procedure followed by fixed-effects inverse-variance weighted meta-analysis (N=3,385). In addition, we conducted a multivariate GWAS of the total plasma N-glycome on groups of N-glycome traits (Supplementary Table 2b) based on the biochemical similarity between glycans measured using MALDI-MS technology. The multivariate analysis was performed using the MANOVA-based method, adopted for analysis of a group of single-trait GWAS summary statistics (the MultiABEL R package)(41). We performed genetic association analyses on 143 univariate (Supplementary Table 2a) and 24 multivariate traits (Supplementary Table 2b).

### Replication

Since we cannot fully match the N-glycan traits measured by UHPLC-FD and MALDI-MS, for each of the 19 glyQTLs not replicated in Sharapov et al. (5), we applied the Cauchy aggregation test (13) to combine P-values across all glycan traits in UHPLC-FD and MALDI-MS N-glycome datasets. P-values were combined using the CCT function of the R package STAAR, which aggregates p values using the Cauchy method. We conducted a one-sided Fisher’s combined probability test for meta-analysis (14) of Cauchy aggregated p-values from UHPLC-FD and MALDI-MS datasets, resulting in replication p-values (Supplementary table 3a). A locus was considered replicated if the Fisher’s p-value exceeded the threshold of 0.05/19=2.63E-03, where 19 is the number of selected glyQTLs.

### Annotation of glyQTLs

We annotated all 59 loci found previously in the study of Sharapov et al. (5) by calculating coefficient of determination R^2^ of association between a locus and two groups of traits – 117 UHPLC-FD-measured N-glycome traits (Supplementary Table 6a) as well as 143 MALDI-MS-measured N-glycome traits (Supplementary Table 6b). Glycan graphical representations following the recommendations of the Consortium for Functional Glycomics (CFG) are presented in Supplementary Tables 2a, 2c, 2d.

### Prioritization of candidate genes in found loci Functional annotation of genetic variants

We inferred the possible molecular consequences of genetic variants in glyQTLs. We focused on variants in LD with lead variants. We created a “long list” of putative causal variants using PLINK version 1.9 (--show-tags option) (42), applied to whole genome re-sequenced data for 503 European ancestry individuals (1000 Genomes phase 3 version 5 data). The size of the window to find the LD was equal to 500 kb. The default r^2^ > 0.8 value was taken as a threshold to include SNPs into the credible set. Prioritization of genes containing variants in strong LD with the lead variant, which are protein truncating variants (annotated by Variant Effect Predictor, VEP (43)) (Supplementary Table 3b) or damaging according to FATHMM XF (44) (Supplementary Table 3c), FATHMM InDel (45) (Supplementary Table 3d).

### Genes of N-glycan biosynthesis and Congenital Disorders of Glycosylation

We searched for the genes encoding glycosyltransferases – enzymes with a known role in N-glycan biosynthesis (46), located in the ±250 Kb-vicinity of the lead SNPs in glyQTLs. Additionally, we prioritized genes with known mutations that cause Congenital Disorders of Glycosylation according to MedGen database (https://www.ncbi.nlm.nih.gov/medgen/76469) located in the vicinity of ±250 kb from the lead SNPs.

### Colocalization with eQTL and pQTL

To find potential pleiotropic effects of glyQTL on gene expression levels in relevant tissues, we applied Summary data-based Mendelian Randomization analysis followed by the Heterogeneity in Dependent Instruments (SMR/HEIDI) (15) on expression of quantitative trait loci (eQTLs) obtained from Westra Blood eQTL collection (47) from peripheral blood; GTEx (version 7) eQTL collection (48) of liver and whole blood; CEDAR eQTL collection (49) from CD19+ B lymphocytes, CD8+ T lymphocytes, CD4+ T lymphocytes, CD14+ monocytes, and CD15+ granulocytes; and on protein quantitative trait loci (pQTLs) using SomaLogic datasets (50,51). The outcome variable was the N-glycome trait with the most significant univariate association. If glyQTLs were only replicated in multivariate analyses, we used summary statistics for the most significantly associated univariate trait as the primary trait in the analysis.

The results of the SMR test were considered statistically significant if P_adj_ < 0.05 (Benjamini-Hochberg adjusted *P*). The significance threshold for HEIDI tests was set at P = 0.05 (P < 0.05 corresponds to the rejection of the pleiotropy hypothesis) (Supplementary Table 6f, 6g).

### DEPICT

Gene prioritization and gene set and tissue/cell type enrichment analyses were performed using the Data-driven Expression Prioritized Integration for Complex Traits framework (DEPICT) (17) applied to GWAMA summary statistics for the samples of European descent (N = 10,172) (5). DEPICT analysis was conducted for SNPs associated with any N-glycosylation trait at P < 5 X 10^-8^/28. The significance threshold for DEPICT analysis was set at False Discovery Rate FDR < 0.20 (Supplementary Table 3e).

## Acknowledgments

None.

## Funding

The authors thank participants and staff of the Diabetes Care System West-Friesland and the DiaGene Study for their cooperation and support. The Hoorn DCS study was supported by a grant from the Foundation for the National Institutes of Health through the Accelerating Medicines Partnership (no. HART17AMP) and the Dutch String of Pearls Initiative. The Hoorn DCS study was supported by the Amsterdam University Medical Center, Location VUmc. The work of A.T., D.M., A.S., S.Sh., Y.S.A. was supported by the Research Program at the Moscow State University (MSU) Institute for Artificial Intelligence.

## Conflict of interest statement

Y.S.A. is a full-time employee of GSK PLC and receives salary and stock options as compensation. All other authors declare no conflicts of interest. Other authors declare no competing financial interests.

## Author contribution

LMtH designed the Hoorn DCS study and contributed to collecting and analyzing the phenotypic and genetic data. PJME contributed to the data collection of the Hoorn DCS study and the content of the paper. RCS analyzed the Hoorn DCS GWAS data.

EJGS is the principal investigator of the DiaGene study. He made the design, performed the coordination, and contributed to the study methodologies. MvH is member of the management team of the DiaGene study. She contributed to the conception, collection, and coordination of the DiaGene study. She had a strong focus on glycomics. AN analyzed the DiaGene GWAS data.

SN and MW contributed towards data acquisition and interpretation.

AT coordinated the work and communications 2023-2025; performed data analysis (quality control analyses, multivariate analysis, post-GWAS, Table 1, Figure 1, ST2, ST4, ST6), wrote the manuscript.

AS performed data analysis (meta-analysis, Cauchy aggregation test, ST3), wrote parts of the manuscript.

DM performed data analysis (quality control analyses, analysis of pleiotropic effects of sialyltransferase loci on diseases and traits, eQTL/pQTL colocalisation, ST5), wrote parts of the manuscript.

NP performed quality control analyses and helped cohort analysts with getting the analyses right.

SSH set up data analysis protocols, coordinated the work and communication 2021-2022, wrote the manuscript.

YSA developed the study concept, provided overall supervision, interpreted results, and wrote the manuscript.

All authors critically assessed the methods and text of this manuscript.

## Abbreviations

TPNG Total plasma N-glycome

T2D Type 2 diabetes

UHPLC-FD Ultra-high performance liquid chromatography-fluorescence detection

MALDI-MS Matrix-assisted laser desorption/ionization-mass spectrometry

QC Quality control

